# Unveiling Hidden Endophytes by Optimising Identification of Endophytic Bacterial Communities from Wild Grassland Plant Roots

**DOI:** 10.64898/2026.02.16.706108

**Authors:** Sobia Ajaz, Manon Longepierre, Elaine Haskins, Joanna Kacprzyk, Tancredi Caruso

**Affiliations:** School of Biology and Environmental Science, University College Dublin, D04 VIW4, Ireland; Sustainable Agroecosystem group (SAE), 8092, ETH Zürich, Switzerland

**Keywords:** Endophytic bacteria, Root sterilisation, amplicon sequencing, Protocol optimisation

## Abstract

Endophytic bacteria are increasingly recognised for their roles in plant health through symbiosis. However, methodological challenges, such as inconsistent root sterilisation, inefficient microbial DNA extraction, and co-amplification of plant organellar DNA, limit accurate characterisation of these communities, especially in wild grassland plants and non model plant in general. To address this, we developed and tested a streamlined protocol for bacterial endophyte detection from wild grassland plant roots, encompassing surface sterilisation of roots, DNA extraction, clamping of plant internal mitochondrial and chloroplast DNA, and 16S rRNA amplicon sequencing. Our approach minimises plant DNA contamination and yields high-quality microbial profiles. The protocol is adaptable and specific to grassland plant species, offering a standardised foundation for endophyte studies in wild and non-model plants.

**Graphical Abstract:** 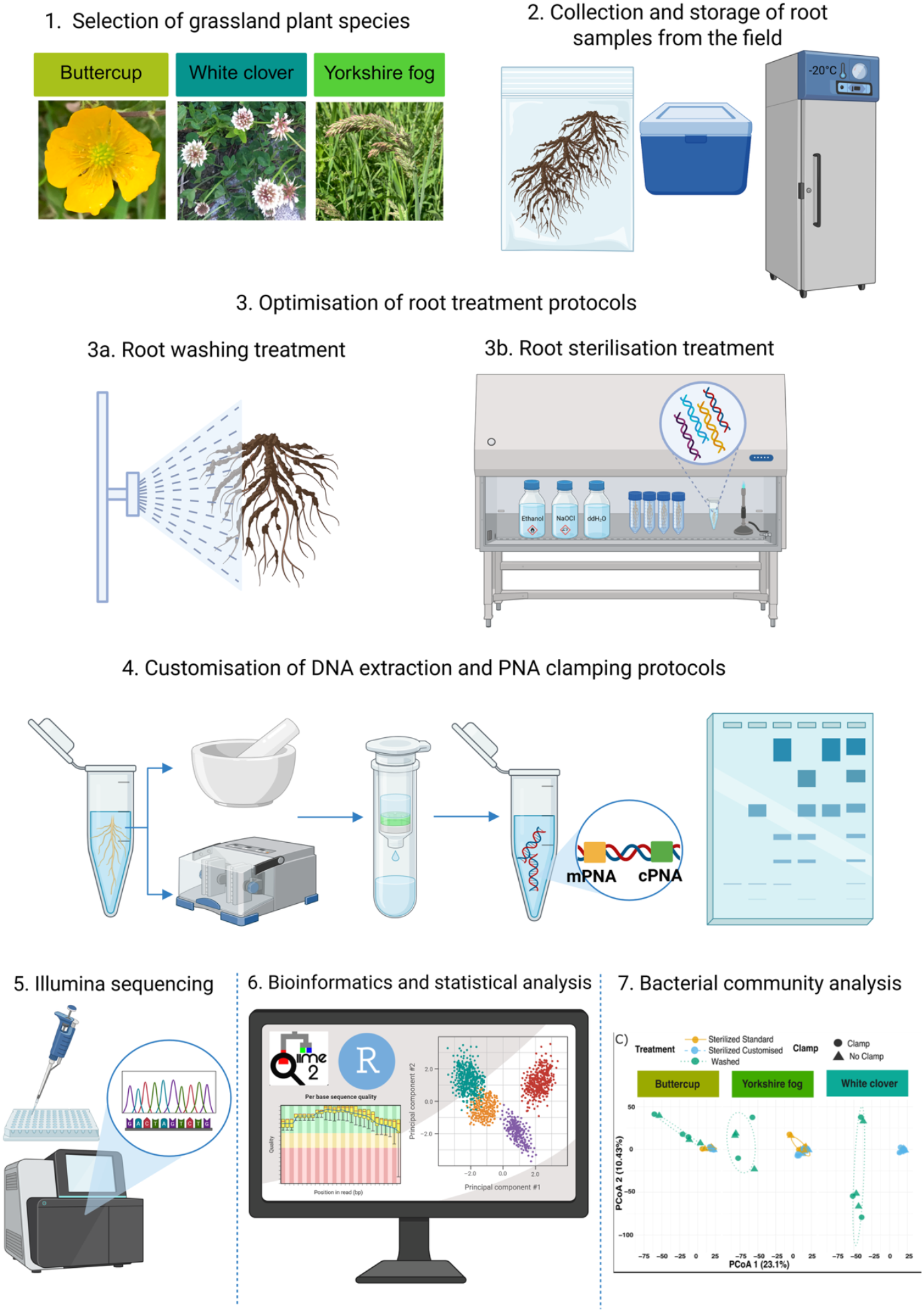

**(Haskins and Ajaz, 2026)** https://BioRender.com/47gd2xr

## Introduction

Endophytic bacteria, residing within plant tissues without causing disease, form complex symbiotic relationships that enhance plant growth, stress tolerance, and disease resistance. They promote plant development through nitrogen fixation (*Azospirillum, Herbaspirillum*) (Guo et al., 2020), phosphate solubilization (*Pseudomonas, Bacillus*) (Lanna-Filho et al., 2022, Berza et al., 2022), and phytohormone production (*Spinacia*) (Carvalho et al., 2014, Chebotar et al., 2023, Misra et al., 2024). Additionally, they suppress pathogens by inducing systemic resistance (*Bacillus subtilis, Streptomyces*) (Safara et al., 2022) and producing antimicrobial compounds (Numan et al., 2022). Their ability to colonise diverse tissues and adapt to environmental stress makes them integral components of plant holobionts, supporting the nutrient cycle, soil health and ecosystem balance. Indeed, their role in plant symbiosis has prompted numerous studies, the results of which can be directly or indirectly applied to the agro-industry (Qadir et al., 2024, Fuentes-Quiroz et al., 2024).

Endophytic detection can be achieved through classical approaches, such as culturing and microscopy, as well as modern genomic techniques using next-generation sequencing, which provide a culture-independent and more comprehensive view of microbial communities (Kaur et al., 2024). The time-consuming nature of microscopy and the fact that ∼90% of microorganisms are unculturable in the lab, make next-generation sequencing one of the most promising approaches for characterising the hidden diversity of endophytic bacteria. Amplicon sequencing, particularly 16S rRNA gene sequencing, has revolutionised the study of endophytic communities by revealing previously undetected microbial richness and providing insights into community composition in various plant species (Berg et al., 2016, Marian et al., 2025). However, challenges remain in distinguishing true endophytes from contaminants or epiphytes due to low biomass, PCR biases, and limitations in resolving closely related taxa, highlighting the need for standardised experimental design and guidelines for data interpretation.

Standardised protocols for isolating endophytic bacteria from plant roots are essential to minimise contamination and ensure reproducibility across studies. A critical step in this process is the surface sterilisation of plant tissues to eliminate external microorganisms (rhizosphere or soil community) without removing internal endophytes (Ding et al., 2019). Commonly, protocols involve sequential treatments with sodium hypochlorite (NaOCl), ethanol, and sterile water, though concentrations and exposure times vary across studies (Wang et al., 2017, Leff et al., 2017, Ben et al., 2017, Robinson et al., 2015, Yang et al., 2021, Guo et al., 2021, Yu et al., 2022). For instance, some methods utilise 5% NaOCl for 5 minutes, followed by rinsing with sterile water, yet residual contamination can still occur (Surette et al., 2003). Alternative approaches, such as sonication (Wei et al., 2018) and ultraviolet (UV) irradiation (Nocker et al., 2007, Górny et al., 2024) have been explored to enhance sterilisation efficacy.

After meticulous surface sterilisation to eliminate external contaminants, the roots undergo a critical mechanical and chemical disruption phase, which is essential for effective DNA extraction and subsequent molecular analyses. The mechanical stage, typically involving techniques such as grinding with a mortar and pestle (often aided by liquid nitrogen) or bead beating, serves to physically break down the robust cell walls of both plant and endophytic bacterial cells (Sambrook and Russell, 2001). This mechanical force liberates the intracellular contents, increasing the accessibility of cellular material to the extraction reagents. Subsequently, the extraction buffer – containing detergents, chaotropic agents, and salts – chemically lyses the remaining intact cells, denatures proteins, and releases high-quality genomic DNA into solution. Together, the combined mechanical disruption and chemical lysis maximise DNA yield and purity, ensuring efficient recovery of both plant and microbial DNA for downstream molecular analyses (Yadav et al., 2018). For instance, molecular techniques like Polymerase Chain Reaction (PCR) rely on adequate template DNA for amplification and subsequent analysis of endophytic microbial communities (Wei et al., 2018). While mechanical disruption is highly effective, careful optimisation is needed to avoid excessive shearing of DNA into smaller fragments, which can affect downstream applications requiring high molecular weight DNA (Yadav et al., 2018).

During PCR, the frequent co-amplification of plant organellar DNA, particularly from chloroplast and mitochondrial sequences, poses a major challenge for studying the bacterial community via 16S rRNA gene metabarcoding (Flörl and Bokulich, 2025, Lundberg et al., 2012). To address this, researchers have developed two primary strategies: (1) designing bacterial-specific primers that minimise binding to plant organellar DNA, and (2) employing PCR clamp inhibition using peptide nucleic acids (PNAs). Bacterial-specific primers, such as 799F (Chelius and Triplett, 2001), reduce chloroplast amplification but may still miss some bacterial taxa, highlighting the need for newer primers with broader coverage (Flörl and Bokulich, 2025). PCR clamp inhibition, particularly using PNAs, has emerged as a powerful tool to block plant DNA amplification by targeting conserved regions in chloroplast and mitochondrial 16S rRNA genes (Lundberg et al., 2012, Sakai and Ikenaga, 2013, Lundberg et al., 2013). These PNA clamps, including PNA-mitochondria and PNA-chloroplast, have been successfully applied in various plant species, such as Arabidopsis, wheat, and citrus (Lundberg et al., 2012, Blaustein et al., 2017, Tkacz et al., 2020). However, their efficacy can be reduced by sequence mismatches in genetically diverse plant species, necessitating customisation for optimal performance (Fitzpatrick et al., 2018a).

To address the challenges of plant DNA interference in 16S rRNA-based studies of endophytic bacterial communities across different grassland species, we conducted a systematic comparison of methodological approaches focusing on *Ranunculus repens* (Buttercup), *Holcus lanatus* (Yorkshire fog grass), and *Trifolium repens* (White clover). This selection provides a methodologically challenging and ecologically relevant testbed: it includes a forb, a grass, and a legume known for its high inhibitor content, thereby stress-testing our protocol across a range of physiological traits. Furthermore, temperate grasslands, where these species are foundational, are vital for agriculture and ecosystem services. Developing a reliable endophyte detection method for this system is therefore directly relevant to understanding plant holobiont function in sustainable agroecosystems.

Our goal was to establish a standardised, reproducible protocol that balances efficiency (maintaining bacterial DNA yield), specificity (minimising plant DNA), and practicality (cost-effectiveness and throughput) suitable for diverse plant species, facilitating more accurate and comparable endophytic microbiome studies across different ecosystems. We hypothesised that optimising root sterilisation and DNA extraction conditions, as well as tailored PNA clamp parameters during 16S rRNA amplification, would (1) significantly reduce plant-derived DNA contamination and (2) decrease sequencing depth differences across grassland species, without compromising bacterial diversity detection.

To test this hypothesis, we systematically evaluated and customised key methodological steps from root sterilisation to PCR amplification. Based on established protocols (Wang et al., 2017, Wemheuer and Wemheuer, 2016), we tested three sterilisation approaches using different chemical concentrations (ethanol/NaClO) and sequential treatments, and evaluated their efficiency in removing surface contaminants while preserving endophytic bacteria.

DNA extraction was performed using commercial Plant and Soil kits, with special attention to overcoming species-specific inhibitors, particularly in White clover, which contains high polyphenol levels (Guo et al., 2021). Subsequently, 16S rRNA gene amplification was performed using bacterial-specific primers (515F-806R) and customised PNA clamp conditions (0.76µM and 2µM concentrations, 15 and 40 sec annealing time of PNA clamp) to suppress chloroplast/mitochondrial co-amplification (Giangacomo et al., 2021, Lundberg et al., 2013) across species. This comparative approach builds on recent methodological advances (Flörl and Bokulich, 2025) while addressing the limitations of universal PNA applications noted in across diverse plant species (Tkacz et al., 2020, Fitzpatrick et al., 2018a).

## Material & Method

### Collection of grassland plant root samples

Three different plant species, *Ranunculus repens* (Buttercup), *Holcus lanatus* (Yorkshire fog grass), and *Trifolium repens* (White clover), were harvested with their roots from six separate agricultural fields in two summer seasons (July 2022 and July 2023) across Ireland. The six distinct sites were respectively located in Lyons (N53.29236,W006.53623), Laois (N52.94324,W007.41193), Longford (N53.73563,W007.95051), Wexford (N52.62762, W006.55917), Galway (N53.309972,W009.14577), and Waterford (N53.40477, W007.93629). White clover was collected from Lyons, Laois, Waterford and Longford; Buttercup was collected from Laois, Waterford, Galway, Longford and Wexford; and Yorkshire fog was collected from Lyons, Laois, Wexford and Galway. These plant species were chosen because they are both abundant and ubiquitous in European grasslands.

Six individual plants of each species were collected at the sites as indicated above. Each set of individual plants was then pooled to create a population-level sample for that site. All plant tissue samples were immediately transferred to 4 °C in the field using a freezer box and moved to a -20°C freezer after collection, where they remained frozen until washed and sterilised.

### Grassland Plant roots samples cleaning process Root washing

The roots were first rinsed roughly under tap water to remove the larger pieces of soil. The roots were then cut with sterile scissors under the safety enclosure. The small root sections were then placed in a clean Falcon tube containing 30-40 mL of epiphyte removal buffer with 0.05 mol/L KH_2_PO_4_, 0.05 mol/L K_2_HPO_4_, and 0.02% Tween 20 (Simmons et al., 2018). Roots and buffer were vortexed (3200 RPM, Voetex-Genie 2) for 2 minutes to remove any attached soil. The rinsing liquid was then discarded and replaced with 30 ml sterilised ultrapure water and 0.01% Tween 20. The sample was vortexed for just 30 s for the second rinse. The wash liquid was again discarded, and 30 ml of ultrapure sterilised water was added for the final rinsing step. The sample was again vortexed for 30 seconds. Finally, the roots were placed in another cleaned Falcon tube using sterile tweezers. These steps were repeated as often as necessary in case the last washing water still contains a lot of soil particles.

### Root sterilisation assay

To test different sterilisation methods, 3 protocols were designed based on different common methods. **Method 1 (M1)** involved sequential washing of roots with 70% ethanol for 2 min, 2% NaOCl for 3 min, and a final 70% ethanol rinse for 2 min. **Method 2 (M2)** included an initial wash with 2% NaOCl for 3 min, followed by 70% ethanol for 2 min. **Method 3 (M3)** consisted of treatment with 0.5% NaOCl for 8 min under constant agitation, followed by 70% ethanol for 2 min. Finally, after all 3 methods, the roots were rinsed twice with ultrapure sterilised water for 3 minutes to remove any chemicals that might interact with the downstream procedure. To test the efficiency of the three methods, 1mL of the last washing water was plated on tryptic soy agar (TSA) medium at 24°C for 48 hours. Immediately after sterilisation, the roots were aseptically placed in a sterile bag using sterile forceps and then instantly frozen in liquid nitrogen. Finally, they were stored for at least 48h at -80°C before DNA extraction.

### DNA extraction assay

The DNA extractions were performed by testing 5 different methods (E1 – E5) derived from the manufacturer’s protocol of the Plant DNA Extraction Kit (DNeasy Plant kit, Qiagen, Germany) and the Soil DNA Extraction Kit (DNeasy Soil kit, Qiagen, Germany). The main protocol modifications were made on the first homogenisation and lysis step for both kits, which combines mechanical and chemical disruption to maximise DNA extraction yield. Root sample preparation was varied, and vortex speed and duration were increased with or without the presence of the CD1 chemical lysing agent buffer supplied with the kits. From the second step onwards, extractions E1, E2, E3 and E4 followed the protocol of the Plant DNA Extraction Kit, while extraction E5 followed the Soil DNA Extraction Kit.

For E1, the protocol performed precisely as outlined by the manufacturer. Briefly, root samples were placed directly into the bead-beating tube containing 800µL of CD1 solution and vortexed twice at 25 Hz with a TissueLyser II (Qiagen, Germany) for 3.5 min. For E2 and E3, root samples were first ground using a sterilised mortar and pestle, then transferred to the bead-beating tube containing 800µL of CD1 solution and vortexed at 30 Hz with a TissueLyser for two 5 min cycles. The only difference between the latter two methods were the types of beads; E2 used the large bead (stainless steel 5mm)supplied in the Dneasy Plant kit, whereas E3 used multiple small zirconium beads provided by the Dneasy Power Soil Pro kit. For E4 and E5, root samples were first lyophilised for 48h with a Freeze Dryer (Labconco™ FreeZone™ Bulk Tray Dryers, 230V Models from Fisher Scientific). Then, the roots were placed in the bead-beating tube without CD1 solution and ground at 30Hz with a TissueLyser for twice 10 min. After 20 min of grinding, the tubes are centrifuged, 800µL of CD1 solution is added, and the samples were vortexed again at 30 Hz for 1 min twice. The major difference between E4 and E5 is that after this first step, the E4 downstream procedure is based on the Plant extraction kit, whereas for E5, it is based on the Soil extraction kit.

Extracted DNA was quantified with Nanodrop2000 (Thermo Fisher Scientific, U.S.A) and standardised to 5ng.µL^-1^ in ultra-pure sterilised water.

### PCR, PNA Clamping and Sequencing

The V4 hypervariable region of the bacterial and archaeal 16S rRNA gene was amplified with the universal primers 515F (5’–GTGYCAGCMGCCGCGGTAA−3’) and 806R (5’-GGACTACNVGGGTWTCTAAT-3’) (Steven et al., 2018). Additionally, two different PNA clamping conditions, mPNA ^5’^GGCAAGTGTTCTTCGGA^3’^ and pNA ^5’^GGCTCAACCCTGGACAG^3’^ (Lundberg et al., 2013) were tested during the amplification in order to evaluate their efficiency.

The first Standard PNA clamping amplification was performed based on (Lundberg et al., 2013) method, using 0.6 µM of each primer with 0.76µM of each PNA clamp with 20ng of standardised template DNA. Prior to use, both PNA clamps were heated to 65°C for 5 min to dissolve all precipitates. The PCR consisted of 38 cycles with an initial denaturation step at 95°C for 40s, followed by a preliminary annealing step for the clamp at 78°C for 15s and a second annealing step for the primers at 58°C for 40s, and finishing with an elongation step at 72°C for 1min.

A Chloroplast PCR assay (ChloPCR) was conducted to optimise the PNA/chloroplast ratio (Supplemental Figure 3a, 3b). For ChloPCR, previously established photosynthetic *Arabidopsis thaliana* (ecotype Col-0) cell suspension culture (Burke et al., 2024) was maintained as described by (May and Leaver, 1993). Cells were grown in full-strength liquid Murashige and Skoog medium (4.3 g /L basal salts, 0.5 mg/L 1-naphthaleneacetic acid, 0.05 mg/L kinetin, 3% w/v sucrose, pH 5.8) in 250 ml Erlenmeyer flasks on orbital shaker (120 rpm), at 22 o C, under 16 h light (50 μmol photons m −2 s −1)/8 h dark regime. All chemicals were purchased from Duchefa, Biochemie (Haarlem, Netherlands). Seven-day-old cells were filtered through Whatman paper (grade 1) using a Buchner funnel under a gentle (water tap) vacuum to remove growth medium before DNA isolation.

DNA extracted from axenic suspension cells of *Arabidopsis thaliana* (ecotype Colombia Col-0) (Burke et al., 2024) was used to test different PCR scenarios on plant samples free of microbial DNA. Three different PNA clamp concentrations were tested: 0.76 µM, as the Standard Clamping Concentration (Steven et al., 2018, Lundberg et al., 2013), as well as 1 µM and 2 µM. Additionally, two different PNA annealing times were tried: one at 15s (Steven et al., 2018, Lundberg et al., 2013) and a second at 40s. Seven chloroplast DNA concentrations (40 ng, 10 ng, 1 ng, 0.1 ng, 0.01 ng, 0.001 ng, and 0.0001 ng) were run under those previously described PCR conditions, resulting in a total of 42 different amplification reactions.

A second PCR assay, termed Customised Clamping, was designed to test whether more stringent conditions could further improve host DNA suppression. Based on the ChloPCR results (Supplemental Figure 3), which indicated that higher PNA concentration and longer annealing times enhanced chloroplast blocking, we increased the PNA concentration to 2 µM and extended the duration of the clamp annealing step to 40s. All other parameters (38 cycles, 0.6µM primers, 20ng DNA) remained identical to the Standard Clamping protocol.

For each PCR assay (Standard Clamping and Customised Clamping), PCR amplification was performed in technical triplicate, and products were pooled before sequencing. The pooled PCR products were sent to the Centre for Genomic Research (Liverpool, United Kingdom) for barcoding and paired-end sequencing using a 2x300bp configuration on the Illumina MiSeq v3 platform (Illumina Inc., San Diego, CA, USA).

### Bioinformatics analysis

QIIME2 version 2024.2 (Bolyen et al., 2019) was used for bioinformatics analysis of the raw 16S rRNA amplicon sequencing data. The raw demultiplexed paired-end sequences were imported into the Qiime2 environment using qiime tools import and Casava 1.8 demultiplexed (paired-end) format. Primer sequences were trimmed within the QIIME2 environment from paired-end reads using the q2-cutadapt plugin (version 1.2.1 (Martin, 2011)). The trim-paired command was run with the forward (--p-front-f) and reverse (--p-front-r) primer sequences. The default error rate of 0.1 was used, allowing for a 10% mismatch rate between the read and primer sequence to account for potential sequencing errors.

DADA2, within QIIME 2, was used to denoise the reads. This step included quality filtering, reads with an average quality score <35. The paired-end reads were joined at a mean length (257± 5.14 bp), with an overlap of 150bp and finally non-overlapping regions and chimaeras were discarded. The sequences were clustered into amplicon sequence variants (ASVs) using the q2-dada2 plugin and the denoise-paired command (Callahan et al., 2016).

A Naïve Bayesian classifier approach, with the feature-classifier plugin and classify-sklearn method in QIIME2, was used to taxonomically assign ASVs from domain to species level using the SILVA database version 138 (Quast et al., 2012). The resulting artefacts of the DADA2 step, including FeatureTable [Frequency] and Feature-Data [Sequence]. For the entire dataset, in total 10,747,070 reads, 111,948 average reads per sample and 16, 638 unique bacterial ASVs were observed. The entire bioinformatics workflow was scripted to ensure reproducibility and transparency, and it’s publicly available at a GitHub repository(https://github.com/sobia-ajaz/Endophytes_wildgrassplants). Raw sequence reads of bacteria have been submitted to the NCBI SRA Database, which is publicly available with accession SAMN54743423-SAMN54743492 as of the manuscript’s acceptance.

Functional annotation of 16S rRNA sequences was conducted through PICRUSt (Phylogenetic Investigation of Communities by Reconstruction of Unobserved States) version 2.5.2. KEGG (Kyoto Encyclopedia of Genes and Genomes) was used to reveal orthologs (Douglas et al. 2020).

### Statistical analysis

Statistical analyses were performed using R version 4.5.0 (2025.04.11). QIIME2’s key outputs, such as feature tables, phylogenetic trees, and taxonomic assignments, were exported and imported into R via qiime2R (v 0.99.6) (https://github.com/jbisanz/qiime2R) and phyloseq (v1.52.0) (McMurdie and Holmes, 2013) for downstream analysis. All samples in the ASV table were rarefied (40,000 reads without replacement, using a fixed random seed to maintain reproducibility) to an even sequencing depth for computations.

The complete ASV table was first used to quantify and visualise the proportion of reads assigned to plant mitochondria, host chloroplasts, and bacterial 16S rRNA genes for each plant species. Statistical comparison was then made between root treatments: sterilised vs washed samples and among PCR procedures: standard vs customised PCR, with or without pPNA + mPNA clamps. These analyses assessed the impact of sample preparation and PCR strategy on the relative abundance of host-derived versus bacterial reads. After that, ASVs related to the host’s mitochondria and chloroplasts were removed from the data for additional analysis on the bacterial community.

Alpha diversity metrics were assessed through the Shannon diversity index, calculated on rarefied bacterial ASV datasets. The significant differences between the three plant species were tested within each of the five treatment groups: Sterilised Standard Clamp, Sterilised Optimised Clamp, Sterilised No Clamp, Washed Clamp, Washed No Clamp, using Kruskal-Wallis tests (Kruskal and Wallis, 1952) followed by pairwise Wilcoxon tests (Wilcoxon, 1945) with Benjamini-Hochberg p-value adjustment (Benjamini and Hochberg, 1995).

The bacterial ASV abundance data were transformed using the centred log-ratio (CLR) transformation to address compositionality (Gloor et al., 2017). Beta diversity was evaluated using principal coordinates analysis (PCoA) based on Euclidean distance of CLR-transformed data. Permutational multivariate analysis (PERMANOVA (Anderson, 2001) was used to test the effects of root preparation methods, plant species, and clamp status on microbial community composition. PERMANOVA was performed using the adonis2 function from the vegan package (Oksanen, 2015) with 999 permutations.

All visualisations were created using ggplot2 (v3.5.2) (Wickham, 2016) with multi-panel layouts to compare patterns across treatment groups.

### PNA Clamp Analysis

Two peptide nucleic acid (PNA) probes targeting plant organellar 16S rRNA regions were tested: a chloroplast-specific probe (5′-GGCTCAACCCTGGACAG-3′) and a mitochondrial-specific probe (5′-GGCAAGTGTTCTTCGGA-3′). For each probe, both forward and reverse complement orientations were aligned against representative chloroplast and mitochondrial ASV sequences using local pairwise alignment in the Biostrings (Pages et al., 2013, Pagès et al., 2025) and pwalign (Aboyoun and Gentleman, 2025) R packages. Alignments were performed with a match score of +1, a mismatch penalty of 1, and gap opening and extension penalties of 1e^6^ to enforce gap-free alignments consistent with the fixed binding length of PNAs. For each ASV, the alignment with the lowest number of mismatches was retained as the best hit.

Organellar ASVs were classified based on their probe alignment mismatches results, defined as: Perfect Match (0), 1 Mismatch, 2 Mismatches, and ≥3 mismatches. These mismatches were merged with their abundance table and experimental metadata. For analysis, Organellar ASV read counts were aggregated across three focal plant species (Buttercup, Yorkshire fog and White clover) and stratified by clamp treatment and PCR protocols (standard or customised). For each condition, the total number of reads per mismatch category was calculated, and relative abundances were expressed as percentages of the total reads. Mismatch distributions were visualised as stacked bar plots using the R package ggplot2 (v3.5.2) (Wickham, 2016).

## Results

### M2 is an excellent Root Sterilisation method

To evaluate the effectiveness of different sterilisation methods in removing epiphytic and rhizoplane bacteria, 1000 µL of the final ultra-pure rinse water from each method (M1, M2, and M3) were plated onto TSA medium and incubated at 24°C for 48 hours. The number of colony-forming units (CFUs) varied across methods (Supplemental Figure 1). M1 colony number ranged from 11 to 156 colonies, giving an average of 29 with a standard deviation of 31; M2 colony number ranged from 1 to 41 colonies, giving an average of 14 with a standard deviation of 10 and M3 colony number ranged from 4 to 177 colonies, giving an average of 30 with a standard deviation of 38. M2 exhibited the lowest average contamination level, suggesting it was the most effective sterilisation method. Statistical analysis, Tukey’s HSD (ANOVA, p < 0.05) confirmed significant differences between M2 and M1/M3, indicating that the protocol using only NaOCl (2% for 3 min) followed by ethanol (70% for 2 min) was more effective than the others.

### E5 is the best DNA extraction with high Bacterial DNA recovery

DNA yield and purity were assessed for the five extraction methods (E1-E5). DNA concentration was measured using a Nanodrop spectrophotometer, and contamination levels were evaluated based on the A260/A280 ratio (Supplemental Figure 2). E5, based on the Soil DNA Extraction Kit (DNeasy Soil kit, Qiagen, Germany), delivered the highest yield and the best purity, likely due to its customised bead-beating and chemical lysis steps. Lyophilisation before extraction (E4 and E5) significantly improved DNA yield compared to fresh root grinding (E1-E3). The difference in yield between E4 and E5 suggests that the Soil DNA extraction Kit’s proprietary reagents better enhanced bacterial DNA recovery in comparison to the Plant DNA extraction kit (DNeasy Plant kit, Qiagen, Germany).

### Host DNA contamination and bacterial DNA recovery differ between sterilised and washed roots

The Raw 16S rRNA amplicon sequencing yielded a total of 15,646,367 paired-end reads. The DADA2 pipeline produced 16,638 bacterial amplicon sequence variations (ASVs) across all samples after merging. Application of root sterilisation or washing techniques significantly altered the bacterial community composition across Buttercup, Yorkshire fog, and White clover. Moreover, the relative abundance of bacterial, chloroplast, and mitochondrial sequences—and their responsiveness to clamping strategies—varied strongly by plant species (Figure 1).

**Figure 1:**
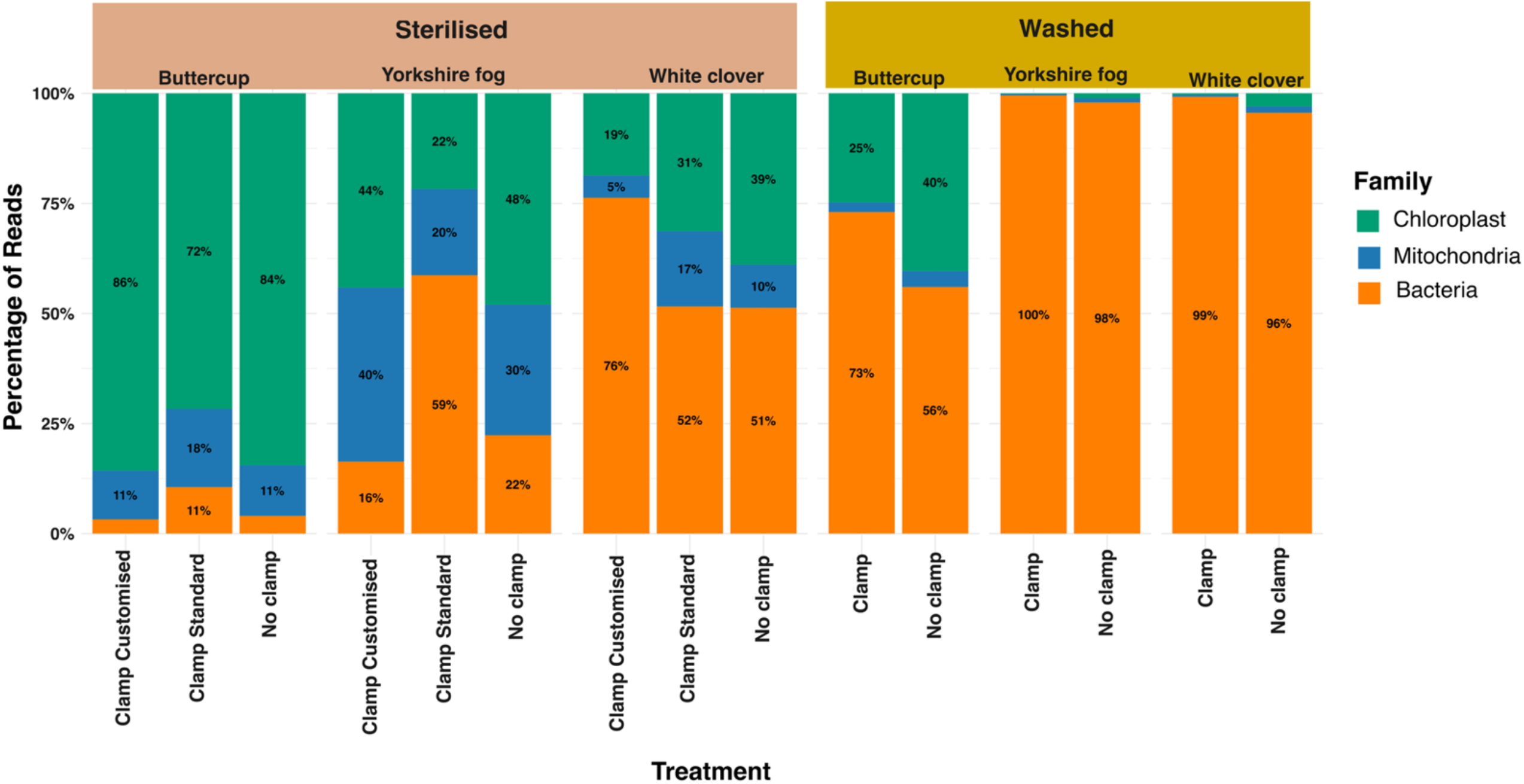
Relative abundance of chloroplast, mitochondrial, and bacterial sequences across plant species and treatments. Stacked bar plots show the proportion of reads assigned to chloroplast (green), mitochondria (blue), and bacteria (orange) for three plant species: Buttercup, Yorkshire fog, and White clover. Samples were processed either as sterilised (left panel) or washed (right panel) roots before DNA extraction. Within each species, different PCR conditions were tested, including Clamp Customised, Clamp Standard, and No Clamp (or Clamp vs No Clamp for washed samples). The y-axis represents the percentage (%) of the reads per treatment, with percentages within bars indicating the relative abundance of sequences for each taxonomic group.

Analysis of host-derived DNA contamination revealed distinct patterns between epiphytic (washed) and endophytic (sterilised) fractions that varied substantially across plant species. In non-clamped samples, sterilised roots exhibited high proportions of host-derived sequences, whereas washed roots were predominantly composed of bacterial reads (Figure 1).

Buttercup (*Ranunculus repens*) exhibited pronounced host DNA contamination, with chloroplast sequences dominating sterilised roots (84–86% of total reads; approximately 115,000–136,000 reads). Mitochondrial reads accounted for 11% (5,800–17,000 reads), while bacterial representation was minimal (11%; 5,000–10,000 reads). In contrast to the sterilisation treatment, washing alone resulted in bacterial reads accounting for a substantially larger portion of reads, with 73–75% of total sequences (94,000–117,000 reads), accompanied by much smaller chloroplast (35,000 reads) and mitochondrial (<2%) contributions.

Yorkshire fog (*Holcus lanatus*) Yorkshire fog exhibited a distinct contamination profile in sterilised samples, characterised by an unusually high proportion of mitochondrial DNA (∼30%; ∼117,000 reads), alongside substantial contributions from chloroplast (∼48%; ∼191,000 reads) and bacterial (∼22%; ∼93,000 reads) sequences.

White clover (*Trifolium repens*) displayed a more balanced distribution in sterilised roots, with chloroplast (39%; 104,000–120,000 reads), mitochondrial (10%; ∼93,000 reads), and bacterial (51%; 91,000–119,000 reads) sequences all represented. Following washing alone, bacterial sequences were predominant (96–99%; 117,000–123,000 reads), while host-derived sequences were reduced to minimal levels (<3,500 reads combined).

### Differential impact of PNA clamping on bacterial DNA recovery

PNA Clamping improved bacterial recovery to varying degrees across plant species within sterilised samples (Figure 1). In Buttercup, clamping modestly reduced chloroplast abundance (136,000 → 115,000 reads) and increased bacterial reads to ∼17,000. In the Yorkshire fog, the Standard Clamp yielded a strong increase in bacterial reads (93,000 → 152,000) while sharply decreasing chloroplast (117,000 → 12,000) and mitochondrial (191,000 → 10,000) reads. The Customised Clamp produced smaller gains (∼67,000 bacterial reads). In White clover, both clamping designs improved bacterial detection: Standard Clamp ∼119,000 bacterial reads; Customised Clamp ∼123,000 bacterial reads with near-complete suppression of mitochondria (< 1,700 reads).

When comparing sterilised and washed roots, washed samples consistently produced the highest bacterial sequencing read abundances across all species. Buttercup bacterial reads rose from ∼17,000 (Sterilised Clamp) to ∼94,000 (Washed Clamp); Yorkshire fog increased from ∼152,000 to ∼270,000; and White clover remained similar (∼119,000 vs ∼123,000), consistent with its higher intrinsic endophytic load.

### PNA Clamp efficiency and mismatch profiles explain differences in host DNA suppression

Mismatch analysis of the chloroplast and mitochondrial PNA clamps revealed strong species-specific differences in sequence complementarity that help explain variation in organellar read suppression (Figure 2). For chloroplast clamps (Figure 2a), both designs achieved high overall binding efficiency, but the Standard Clamp outperformed the Customised Clamp in Buttercup (92.5 vs 87%) and White clover (97.4 vs 79.4%). In contrast, Yorkshire fog showed better compatibility with the Customised Clamp (81.5% perfect matches) than with the Standard Clamp (57.4%). Contrastingly, mismatched reads (one–two mismatches) reached > 40%. For mitochondrial clamps (Figure 2b), perfect matches were frequent in Buttercup (80.6–91.3%) but much lower in Yorkshire fog (49–72%) and White clover (8–18%). In these latter species, mismatched sequences dominated (up to 60% with two mismatches).

**Figure 2:**
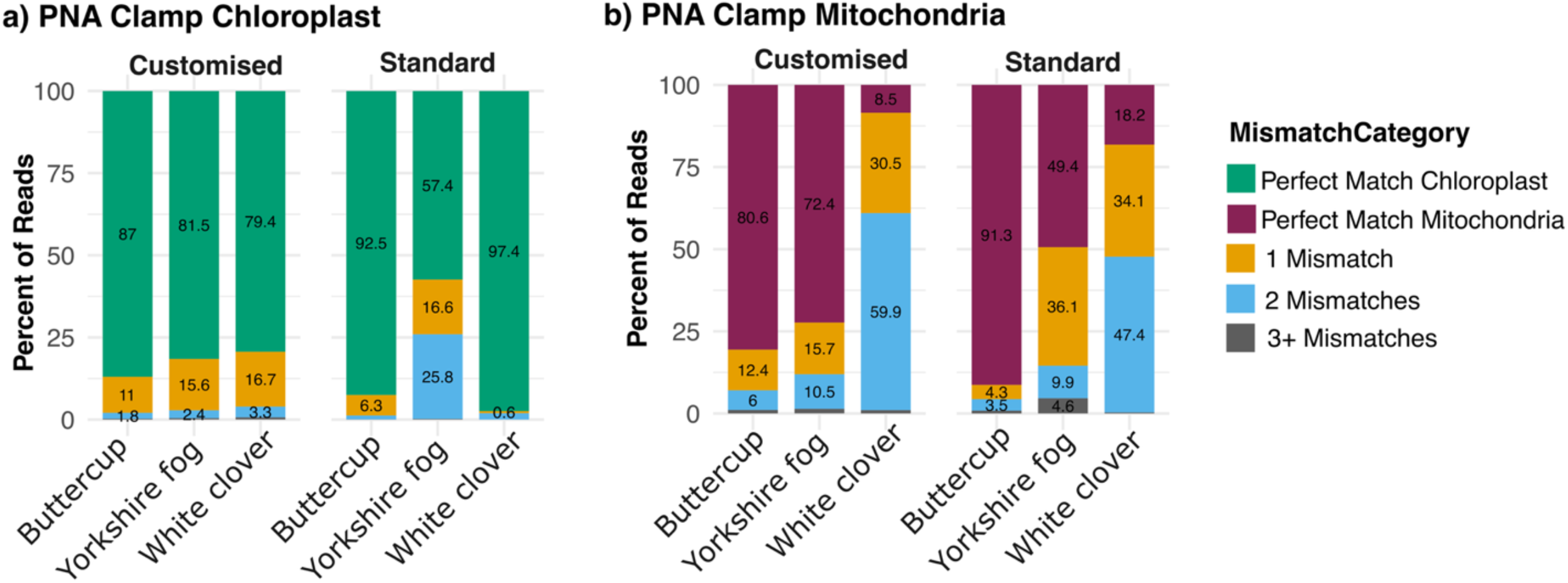
Suppression of chloroplast (a) and mitochondrial (b) reads by PNA clamps across plant species. The proportion of reads is shown for three plant species (Buttercup, Yorkshire fog, and White clover) under different clamp treatments (no clamp, standard clamp, and customised clamp). Bars are subdivided by mismatch categories (perfect match, 1 mismatch, 2 mismatches, 3+ mismatches), representing the degree of binding between the PNA clamp and chloroplast/mitochondrial templates

Comparison of mismatch profiles (Figure 2) with organellar read suppression patterns (Figure 1) revealed that predicted PNA–target complementarity did not consistently correspond to proportional increases in bacterial read abundance across species. While improved matching aligned with enhanced host DNA suppression in Buttercup, this relationship was less direct in Yorkshire fog and White clover, indicating that sequence complementarity alone does not fully predict in-sample clamp performance.

### Host identity, clamping, and sterilisation jointly structure root-associated bacterial communities

Bacterial community composition and diversity varied strongly among host species and treatments (Figure 3A–C). Sterilised roots, representing the endophytic fraction, were less diverse and typically dominated by one or two bacterial families, whereas washed roots, encompassing both epiphytic and endophytic bacteria, supported richer and more even communities. Across all comparisons, host species exerted the greatest influence, producing clear, species-specific bacterial profiles in *Ranunculus repens (*Buttercup), *Holcus lanatus* (Yorkshire fog), and *Trifolium repens* (White clover).

**Figure 3:**
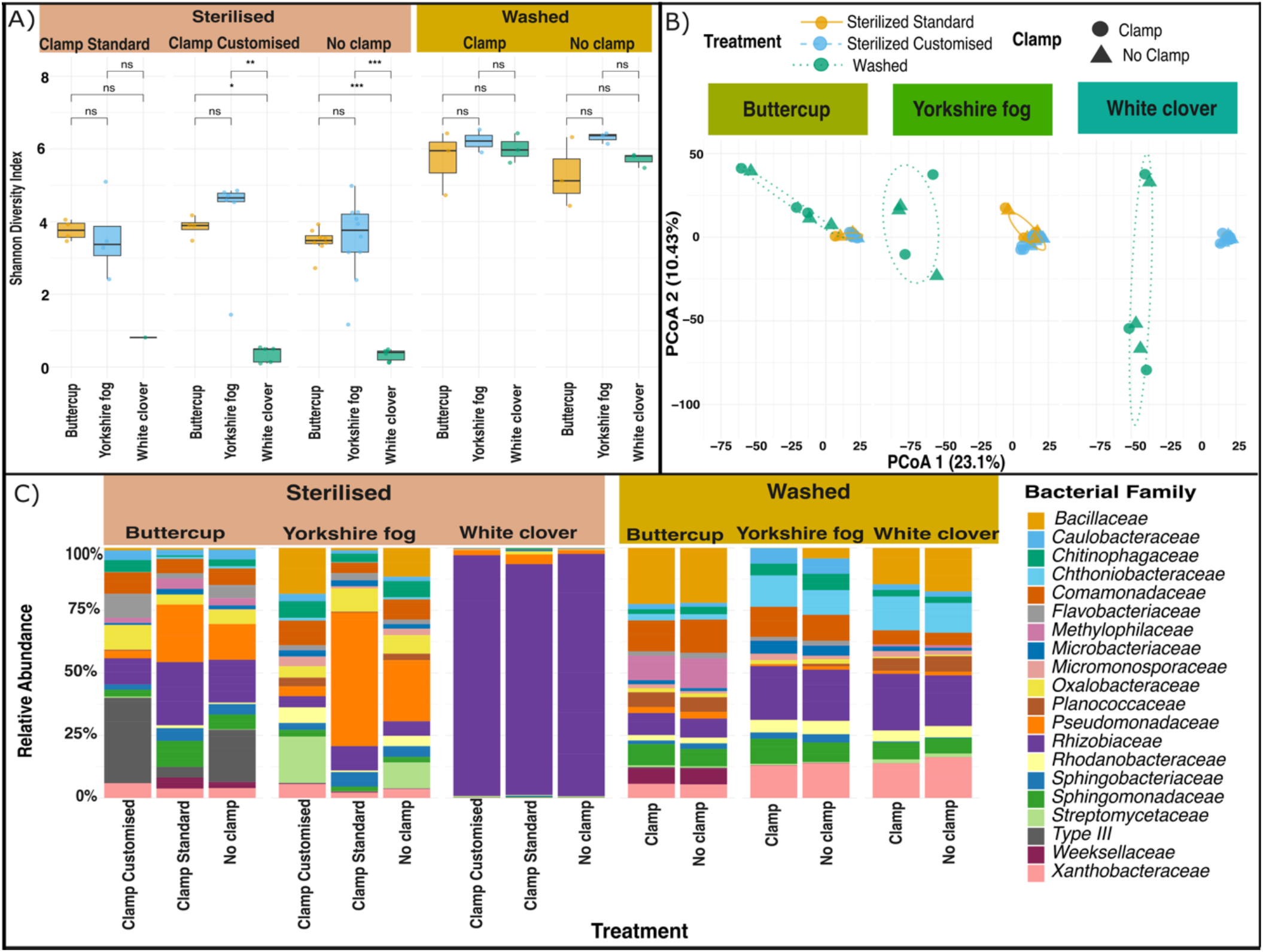
Bacterial community analysis of plant microbiomes across different treatments. A) Alpha diversity analysis using the Shannon index. Box plots display the Shannon diversity index for each of the above plant species under different root preparation treatments and PCR conditions. Significant differences between groups, calculated using Kruskal-Wallis tests, are indicated by asterisks (p<0.05, p<0.01, p<0.001) and “ns” denotes no significant difference. B) Multivariate analysis of bacterial community structure across plant species and treatments. The Principal Coordinates Analysis (PCoA) is based on centred log-ratio (CLR)–transformed Euclidean distances. Each panel represents a different plant species. Colours denote treatments (Sterilised Standard, Sterilised Optimised, and Washed) and symbols indicate Clamp status (Clamp vs. No Clamp). The percentage of total variance explained by each axis is shown in parentheses. C) Taxonomic composition of bacterial families. Bar plots show the relative abundance (%) of the twenty most abundant bacterial families detected in samples from three different wild grassland plant species, i.e. Buttercup (*Ranunculus repens*), Yorkshire fog (*Holcus lanatus*), and White clover (*Trifolium repens*). Samples are categorised by treatment (Sterilised or Washed) and by PNA clamping method (Customised or Standard). The colour-coded legend to the right indicates the bacterial family corresponding to each bar segment

Alpha diversity, quantified using the Shannon index—which reflects both richness and evenness—differed significantly among host species and treatments (Figure 3A). The strongest effect was observed between root preparation methods: washed roots consistently showed higher bacterial diversity than sterilised roots. Under sterilisation, both PNA clamping procedures (Standard and Customised) increased bacterial diversity across all three plant species, whereas this difference disappeared when roots were only washed. Although mean alpha diversity did not clearly differ between the two clamping methods, the customised clamp tended to yield lower variability among samples. At the plant-species level, Buttercup and Yorkshire fog exhibited higher diversity than White clover under sterilisation, whereas differences among species were less pronounced under washed conditions. Overall, the root cleaning method was the dominant driver of alpha diversity, followed by the plant species, with the clamping procedure having a host-dependent effect.

The Principal Coordinates Analysis (PCoA) revealed strong clustering by host species and clear separation between sterilised and washed samples (PCoA 1 = 23.1 %, PCoA 2 = 10.4 %) at the bacterial community level. Buttercup, Yorkshire fog, and White clover formed distinct, non-overlapping clusters. Within every plant species, sterilised and washed samples were separated primarily along the first axis. These ordination patterns were supported by PERMANOVA, which showed that treatment was the strongest driver of bacterial community composition (*R*^*2*^ *= 0*.*22, F = 8*.*56, p = 0*.*001*), while host species also had a significant but smaller effect (*R*^*2*^ *= 0*.*07, F = 2*.*23, p = 0*.*004*) (Supplementary Table 1). The influence of clamping was most pronounced within sterilised roots: both clamp types shifted Buttercup and Yorkshire fog communities away from the no-clamp clusters, whereas in White clover, clamping produced minimal community structural shift. In washed samples, clamped and unclamped replicates overlapped extensively. Consistent with these patterns, clamp status did not significantly influence overall bacterial community composition (PERMANOVA, *R*^*2*^ *= 0*.*006, F = 0*.*35, p = 1*.*00*) (Supplementary Table 1). Overall, these ordination patterns demonstrate that beta diversity was structured primarily by host species and sterilisation, with clamping exerting a secondary, host-dependent effect.

When looking at the bacterial community composition assigned at the family level, sterilised and washed samples showed very different taxonomic patterns, and each plant species harboured a distinct microbial community. In Buttercup, sterilised roots were dominated by *Rhizobiaceae, Pseudomonadaceae*, and unclassified (“Type III”) families, forming a consistent core across clamping treatments. Minor contributions came from low-abundance *Comamonadaceae, Caulobacteraceae, Oxalobacteraceae, Rhodanobacteraceae, Sphingomonadaceae, Xanthobacteraceae* and *Microbacteriaceae*. The Customised clamp produced a slight reduction in *Rhizobiaceae* and a corresponding increase in *Comamonadaceae, Flavobacteriaceae, Oxalobacteraceae* and unclassified (“Type III”) families. In contrast, the Standard clamp maintained high *Rhizobiaceae* dominance but increased the relative abundance of *Shingomonadaceae* and *Weeksellaceae*. However, in the roots of washed Buttercup, higher taxonomic richness was found. In Buttercup washed roots, *Bacillaceae, Planococcaceae* were found in high abundance, while they were not detectable in the case of sterilization. In the washed treatment, clamping effects were less pronounced.

In the Yorkshire fog, sterilised unclamped samples were dominated by *Pseudomonadaceae, Streptomycetaceae* and *Bacillaceae* with additional contributions from *Chitinophagaceae, Comamonadaceae, Oxalobacteraceae, Rhizobiaceae, Rhodanobacteraceae* and *Sphingomonadaceae*. The taxonomic composition of the less abundant ones was different depending on thethe clamping strategy. The standard clamping strongly suppressed *Bacillaceae* and enriched *Pseudomonadaceae, Rhizobiaceae, Sphingomonadaceae and Oxalobacteracea*. Contrastingly, customised clamping increases the relative abundance of *Bacillaceae* and *Streptomycetaceae* while reducing *Pseudomonadaceae, Rhizobiaceae species*. Washed roots in the *Yorkshire fog* hosted the most compositionally even community. The six dominant families were *Rhizobiaceae, Comamonadaceae, Sphingomonadaceae, Caulobacteraceae, Chthoniobacteraceae*, and *Xanthobacteraceae*. Clamping in washed samples suppressed *Bacillaceae* and caused only a minor shift in the abundance of several other families.

In White clover, sterilised roots were almost exclusively composed of *Rhizobiaceae* (> 90 %) across all clamping treatments, with only a small contribution of *Pseudomonadaceae* (< 5 %). Clamping exerted minimal influence on community composition, although Standard clamping allowed the detection of very low-abundant families such as *Oxalobacteraceae* and *Microbacteriaceae*. Washed clover roots showed a markedly more diverse community composition. *Rhizobiaceae* remained dominant (20-25%) but exhibited an increased representation of *Bacillaceae, Chthoniobacteraceae, Comamonadaceae, Planococcaceae, Rhodanobacteraceae, Sphingomonadaceae* and *Xanthobacteraceae*. These families were not detected in sterilised roots. Clamping in the washed roots does not change the bacterial taxonomic structure, but causes minor changes in the relative abundance of certain families.

The functional prediction data (e.g., transport systems, transcriptional regulators, sigma factors, and chemotaxis proteins) also highlight that sterilised root treatments yielded a higher abundance of predicted KEGG functions compared to washed-only controls (Supplemental Figure 7). This indicates that surface contaminants can mask the true diversity and functionality of endophytes. By applying sterilisation, the sequencing effort captures more endophytic-specific reads, leading to improved recovery of functional genes such as ABC transporters, amino acid transport systems, and transcriptional regulators.

In Buttercup roots, sterilised samples showed a much higher abundance of predicted KEGG functions compared to washed treatments. Functional categories such as ABC transport systems (sugar, peptide/nickel, iron complex), transcriptional regulators, and RNA polymerase sigma factors were consistently more represented in sterilised roots. For Yorkshire fog, the effect of sterilisation was particularly pronounced. Sterilised roots yielded substantially higher quantities of chemotaxis proteins, two-component response regulators, and sugar/peptide transporters, pointing to a rich endophytic bacterial community with active signalling and nutrient exchange functions. Washed-only treatments showed lower functional recovery. In White clover, sterilised root samples again revealed enhanced functional diversity compared to washed roots. Key endophyte-associated functions such as polar amino acid transport systems, ABC transporters, transcriptional regulators, and chemotaxis proteins were significantly enriched in sterilised treatments.

## Discussion

This study systematically deconstructs the methodological pipeline for endophytic microbiome research, revealing how each step, from root surface sterilisation and DNA extraction to the application of PNA clamps, profoundly influences the apparent composition and diversity of bacterial communities. Our findings underscore a central paradox in plant microbiome studies: the very methods designed to reveal hidden microbial players can also distort their portrait. By integrating physical and molecular strategies, we demonstrate that a tailored, multi-faceted approach is paramount for accurately profiling the endophytic niches of wild grassland plants, Buttercup, Yorkshire fog, and White clover, which are often characterised by low microbial biomass and high host DNA contamination.

### Surface sterilisation: A non-neutral filter defining the endosphere

Surface sterilisation is a critical step for distinguishing true endophytes from epiphytic contaminants (Mercado-Blanco, 2014). Our results identify the sequential NaOCl–ethanol protocol (M2) as the most effective, minimising culturable epiphytes in rinse water while presenting endophytic communities. This aligns with literature advocating for sequential chemical treatments to dismantle complex soil biofilms without excessive penetration into the apoplast (Wang et al., 2017, Ding et al., 2019). The failure of less stringent (M1) or prolonged (M3) sterilisation protocols was evident in significantly higher epiphytic CFUs.

The profound impact of sterilisation was confirmed molecularly. Sterilised roots exhibited constant lower alpha diversity than washed roots, and the PCoA revealed a clear separation in community structure, demonstrating that sterilisation fundamentally shifts the observed profile away from the total root-associated microbiota. Taxonomically, washed roots were enriched for classic rhizosphere inhabitants—such as the *Flavobacteriaceae, Comamonadaceae*, and *Bacillaceae* (Edwards et al., 2015, Bulgarelli et al., 2012), whereas sterilised roots exhibited a conserved and host-specific endophytic signature. *Rhizobiaceae* and *Pseudomonadaceae* in Buttercup, *Pseudomonadaceae* and *Streptomycetaceae*, in Yorkshire fog and a dominant *Rhizobiaceae*. signature in White clover. These patterns reflect the strong selective filter of the plant interior (Hardoim et al., 2015).

Thus, the ideal protocol represents a compromise between rigour and practicality. It must be stringent enough to eliminate surface contamination yet gentle enough to avoid damaging endophytes, while also being efficient in time and processing load for realistic experimental workflows. Our comparative evaluation provides an empirical basis for this balance, defining a protocol (M2) that maximises epiphyte removal, quantifies its selective impact, and is pragmatically sustainable. Finally, the functional predictions (PICRUSt2) indicated that effective sterilisation clarifies the endophytic functional signal, yielding a higher relative abundance of genes linked to nutrient transport, chemotaxis, and regulation. Without this step, the functional potential of the true endophytic consortium is obscured by epiphytic noise. This remains true despite the limitations approches such as PICRUSt2 as applied to Amplicon sequencing data: in our case, the main point is we collected evidence that effective sterilization is vital to avoid a functional descriptions biased towards the root biome, and so obscuring the specificity of the strictly endophytic assemblage.

### Mitigating Host contamination: The double-edged sword of efficient lysis and PNA Clamps

The DNA extraction step is a critical determinant of downstream outcomes, influencing not only yield but also the relative representation of plant versus microbial DNA. The superior performance of the lyophilisation-bead beating–PowerSoil protocol (E5) can be attributed to its synergistic combination of physical and chemical lysis. Lyophilisation (freeze-drying) embrittles plant cell walls, making them more susceptible to mechanical disruption, while vigorous bead beating ensures the liberation of endophytes from within tough root tissues (Yadav et al., 2018, Giangacomo et al., 2021). This is particularly vital for Gram-positive bacteria with robust cell walls, which can be underestimated in milder extraction protocols (Vishnivetskaya et al., 2014). Paradoxically, this efficiency creates a new challenge: the simultaneous release of vast quantities of host organellar DNA.

In this context, PNA clamping offers a powerful solution to this problem by selectively suppressing host 16S-like organellar sequences. Our results confirm that chloroplast-targeted PNA clamps are a highly reliable tool, consistently blocking a dominant source of contamination across diverse plant hosts (Lundberg et al., 2013). However, the variable and often poor performance of mitochondrial clamps constrains their universal application. Mismatch analysis suggests that the efficacy of mitochondrial clams depends on complementarity between the PNA probe and the host’s organellar DNA target sequence. This divergence likely reflects the combined influence of organellar copy number, mismatch position, and PCR dynamics, factors that are not captured by sequence-based predictions alone (Lundberg et al., 2013). The high degree of sequence conservation in chloroplast 16S rRNA genes across plants makes them a robust target for a universal clamp (Fitzpatrick et al., 2018b). In contrast, mitochondrial genomes evolve more rapidly, leading to sequence polymorphisms that can introduce mismatches and drastically reduce clamping efficiency (Fitzpatrick et al., 2018b).

Our results also demonstrate that the effectiveness of PNA clamping is not universal and that the most appropriate protocol must be customised to the host species. While our Customised Clamp (2 µM, 40s annealing) was designed based on its superior ability to suppress a pure chloroplast target in the ChloPCR assay, it did not universally improve bacterial community profiling across all plant species tested. In fact, it improved bacterial recovery for White Clover but was detrimental in Buttercup and Yorkshire Fog. Our findings in White clover are a textbook example of this phenomenon, directly supporting the argument made by Fitzpatrick and colleagues for the development of host-specific PNA clamps for genetically diverse species (Fitzpatrick et al., 2018a). Likewise, our observation that increasing PNA concentration and annealing time could partially improve performance for some plant species (e.g., White clover) but was counterproductive in others (e.g., Buttercup) reinforces the need for empirical, species-specific optimisation (Marian et al., 2025).

These findings highlight a critical methodological principle: the goal is not to maximise host DNA suppression at any cost, but to find a species-specific balance that preserves the integrity of the bacterial community signal. We hypothesise that at higher concentrations, PNAs may non-specifically interfere with the amplification of bacterial 16S rRNA genes for some plant communities, potentially through off-target binding to bacterial templates (Gohl et al., 2016) or by generally inhibiting polymerase processivity. This effect appears to be community-dependent, influenced by the unique taxonomic composition of each plant’s microbiome. Crucially, our study adds a significant layer of evidence that PNA clamping acts as a “silent” enhancer: it dramatically improves bacterial sequencing depth when compared to no clamping procedure, allowing the detection of less abundant to rare taxa. This enhanced depth provides the statistical power necessary not only for more robust quantification of common members but, crucially, for reliably detecting and distinguishing these less abundant to rare taxa (Hussain et al., 2025, Taerum et al., 2020).

### Host species as the ultimate determinant of endophytic community structure

Our methodological refinements ultimately serve to clarify the primary biological driver in this system: host species identity. The endophytic communities of Buttercup, Yorkshire Fog, and White Clover were fundamentally distinct, a divergence reflecting deep-seated differences in host phylogeny and physiology. While the soil microbiome provides the regional species pool, our results demonstrate that host-specific selection acts as the decisive final filter within the endophytic niche. This is powerfully illustrated by the contrast between the grass and the forbe which supported more phylogenetically diverse, generalist communities, and the legume White Clover, which harboured a specialised consortium dominated by *Rhizobiaceae*. This pattern is a direct result of the legume’s unique physiology and active recruitment of nitrogen-fixing symbionts, providing a clear empirical demonstration of how host identity overrides environmental gradients to structure the endosphere, aligning with core tenets of microbiome assembly (Hakim et al., 2021, Bai et al., 2015, Fitzpatrick et al., 2018a). The dominant bacterial families identified constitute a coherent functional core, encompassing taxa responsible for mutualistic growth promotion (e.g., *Rhizobiaceae, Pseudomonadaceae*), bioprotection (e.g., *Streptomycetaceae, Bacillaceae*), nutrient cycling (e.g., *Sphingomonadaceae, Comamonadaceae*), and root colonisation (e.g., *Caulobacteraceae*). Collectively, this assembly captures the essential microbial functions that underpin plant health in grassland ecosystems. By employing a stringent pipeline to minimise methodological artefacts, our study provides robust evidence for a host-specific functional assemblage in wild plants, advancing from cataloguing membership to elucidating the ecological basis of plant-microbe interactions.

### Conclusions and Future Directions

In conclusion, there is no single best method but rather a best-fit approach that must be tailored to the host species and research question. Looking forward, a critical next step will have to involve developing a phylogenetically informed toolkit of mitochondrial PNA clamps tailored to common plant families and genera, which would directly address the specificity limitations observed in our study. Furthermore, incorporating synthetic microbial communities spiked into plant tissue extracts could enable a quantitative assessment of biases introduced during both sterilisation and DNA extraction processes, moving beyond qualitative comparisons to provide robust, measurable quality controls. Most significantly, applying this customised methodological framework in tandem with meta-transcriptomic or metabolomic analyses will be essential to transition from merely cataloguing microbial membership to truly understanding functional dynamics, thereby unlocking the physiological secrets of plant-endophyte interactions. By rigorously validating each step of the analytical pipeline, our study offers a foundation for future investigations aimed at exploring plant endophytes with greater accuracy, reproducibility, and ecological relevance.

## Acknowledgments

This study was supported by a Science Foundation of Ireland grant 20/FFP-P/8584. Authors like to thank the farmers in Lyons, Laois, Longford, Wexford, Galway, and Waterford for their generous cooperation, field access, and assistance with sample collection. We would also like to sincerely thank Patrick Doran for his invaluable support during fieldwork and for accompanying us in identifying suitable sites for this study. Their support was invaluable to the successful completion of this study.

## Author contributions

SA, ML and TC conceptualised and designed the manuscript. SA and ML gathered the literature. ML and EH collected the root samples from the field. SA and ML did the molecular work. SA performed bioinformatics analysis, organised and structured the information and wrote the manuscript with ML and TC. JK perform pure culturing of chloroplasts from Arabidopsis cells (ecotype Colombia Col-0). All authors reviewed and contributed to the text of the manuscript.

## Data availability

The sequencing data presented in this study are already submitted to NCBI Bioproject PRJNA1405508 as a plant metagenome with BioSample accessions SAMN54743423-SAMN54743492. It will be publicly available with the acceptance of the manuscript and will be available at (https://www.ncbi.nlm.nih.gov/biosample/54743423). The entire bioinformatics workflow was scripted to ensure reproducibility and transparency, and it’s publicly available at a GitHub repository (https://github.com/sobia-ajaz/Endophytes_wildgrassplants).

## Supporting Information

Supplementary Information S1

