## Supplementary Material S1 for "Unveiling Hidden Endophytes by Optimising Identification of Endophytic Bacterial Communities from Wild Grassland Plant Roots"

**Root Sterilisation** **Methods**

Sterilisation methods

|  | number of colonies in per µL from the second washing Water |
| --- | --- |
| M1 | 29.3182 |
| M2 | 13.7727 |
| M3 | 30.0909 |

Figure 1: Comparative analysis of root sterilisation methods

**DNA Extraction Methods**

DNA Extraction methods

DNA concentration (in ng.µL-1)

| Plant biomass (in g) | E1 |
| --- | --- |
| 0.214 | 28.5 |
| 0.186 | 5.4 |
| 0.128 | 6.2 |
| 0.117 | 5 |
| 0.138 | 6.1 |
| 0.168 | 1 |

| Plant biomass (in g) | E2 |
| --- | --- |
| 0.14 | 31.4 |
| 0.123 | 15.1 |
| 0.118 | 32.2 |
| 0.102 | 20.6 |
| 0.1 | 19.6 |
| 0.059 | 5.8 |

| Plant biomass (in g) | E3 |
| --- | --- |
| 0.112 | 13 |
| 0.115 | 7.5 |
| 0.141 | 10.2 |
| 0.12 | 8.6 |
| 0.126 | 8 |
| 0.065 | 2.4 |

| Plant biomass (in g) | E4 |
| --- | --- |
| 0.018 | 2.9 |
| 0.043 | 4.1 |
| 0.04 | 6.3 |
| 0.041 | 23.8 |
| 0.069 | 28.3 |
| 0.016 | 2.4 |

| Plant biomass (in g) | E5 |
| --- | --- |
| 0.037 | 8.4 |
| 0.034 | 5.6 |
| 0.041 | 13.5 |
| 0.06 | 66.5 |
| 0.027 | 4.4 |

Supplemental Figure 2: Comparative analysis of DNA extraction protocols

**ChloPCR assay**

A.1

A.2

A.3

1	2	3	4	5	6 7 neg

1	2	3	4	5	6	7 neg

1	2	3	4	5	6	7 neg

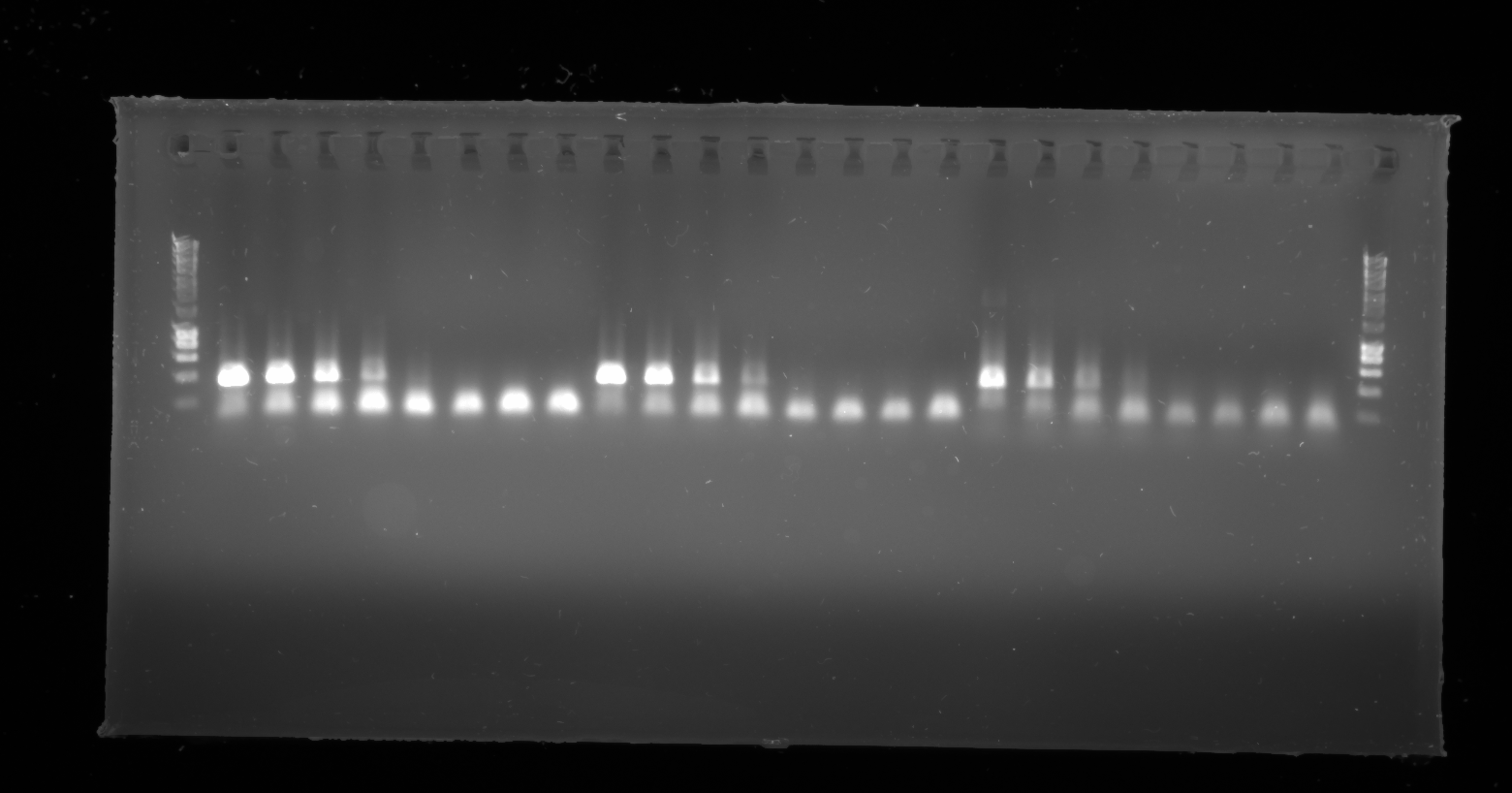

Supplemental Figure 3 (a) The annealing temperature for the PNA clamp in the A conditions was at 15s. The concentration of chloroplast varied across the wells (1-7): 40, 10, 1, 0.1, 0.01, 0.001 and 0.0001ng per reaction and finished with a negative control. The PNA clamp concentrations were 0.76, 1 and 2 µM in A1, A2 and A3, respectively.

B.1

B.2

B.3

1	2	3	4	5	6 7 neg

1	2	3	4	5	6 7 neg

1	2	3	4	5	6	7 neg

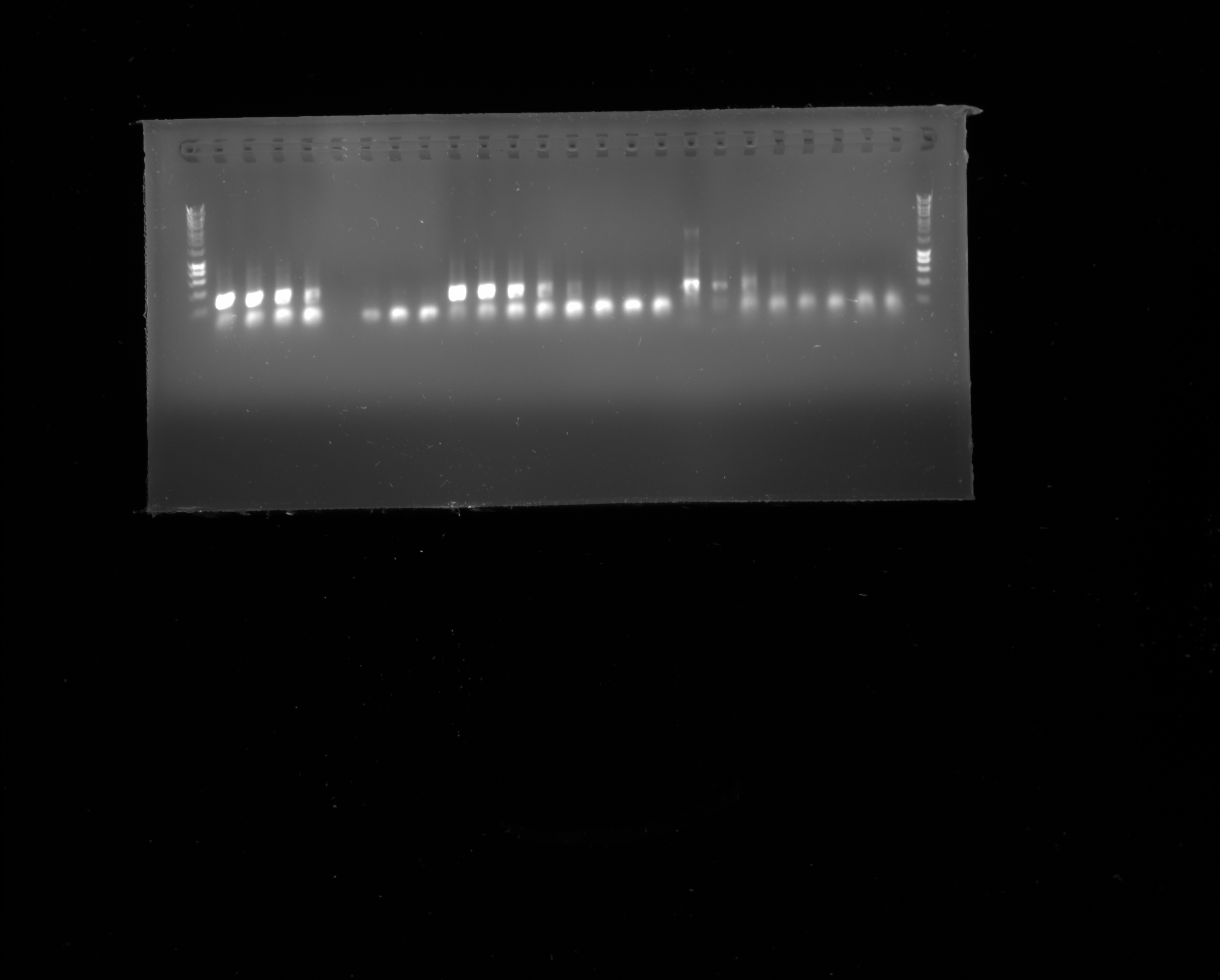

Supplemental Figure 3(B): The annealing temperature for the PNA clamp in the B conditions was at 40s. The concentration of chloroplast varied across the wells (1-7): 40, 10, 1, 0.1, 0.01, 0.001 and 0.0001ng per reaction and finished with a negative control. The PNA clamp concentrations were 0.76, 1 and 2 µM in B1, B2 and B3, respectively.

At 0.76 µM and 1 µM of PNA (Figure A1 - A2 and B1 - B2) for both PNA annealing time (15s Figure A and 40s Figure B), clear bands could be seen at 40, 10 and 1ng of chloroplast in the PCR reaction. For those same conditions but at 0.1ng of chloroplast in the PCR reaction, the bands are very thin. At 2 µM of PNA (Figure A3 and B3) for both PNA annealing time (15s Figure A and 40s Figure B), clear bands could be seen at 40 and 10ng of chloroplast in the PCR reaction. At 1ng of chloroplast in the PCR reaction, the bands are very thin, whereas at 0.1ng of chloroplast in the PCR reaction, the bands are almost non-existent, even more true for the PNA annealing time of 40s.

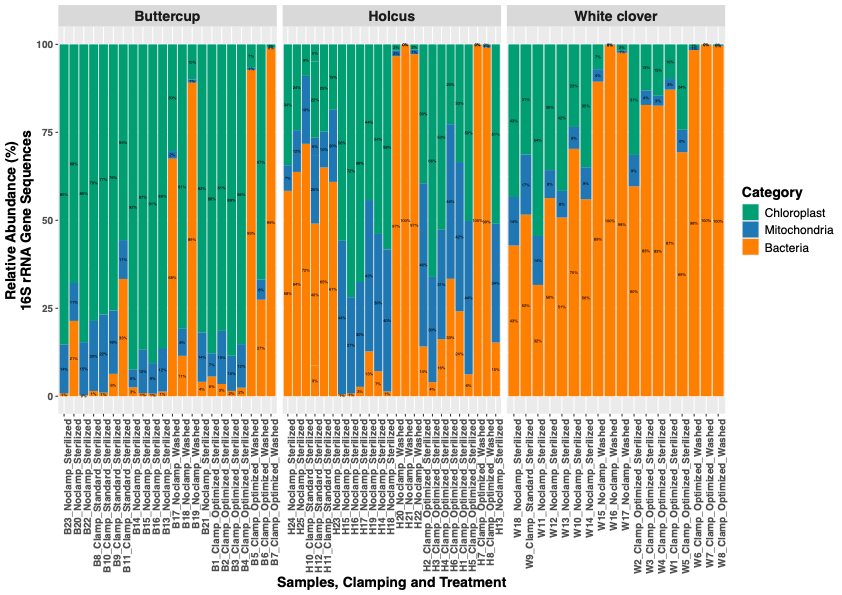

Supplemental Figure 4: Impact of PNA clamping on microbial and host organellar 16S rRNA gene sequence recovery across three plant species. The stacked bar plots represent the relative abundance of bacterial, chloroplast, and mitochondrial 16S rRNA gene sequences from Buttercup, Holcus, and White clover, across different clamping strategies: no clamp, standard PNA clamp, and optimised PNA clamp.

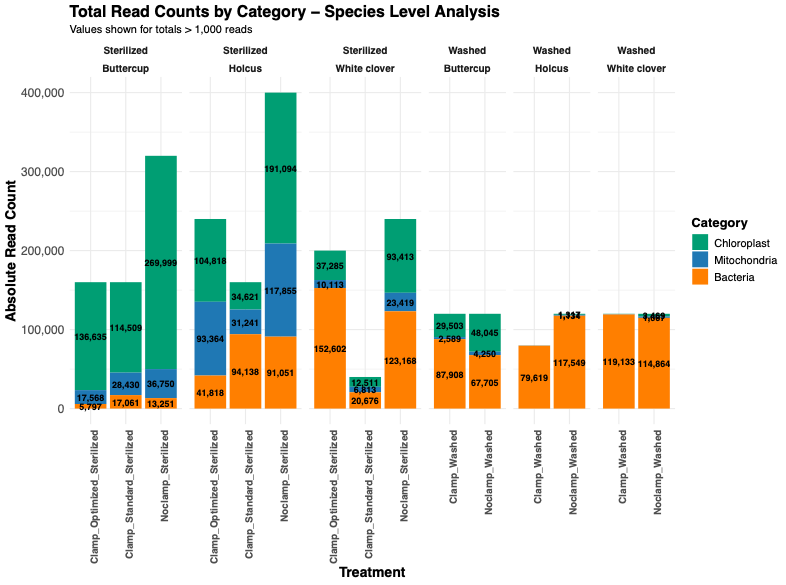

Supplemental Figure 5: Absolute read counts of bacterial, chloroplast, and mitochondrial 16S rRNA gene sequences across clamping treatments in three plant species. Bar plots display the absolute abundance of 16S rRNA gene reads assigned to bacterial families, chloroplast, and mitochondrial sequences across different treatments in Buttercup, Holcus, and White clover.

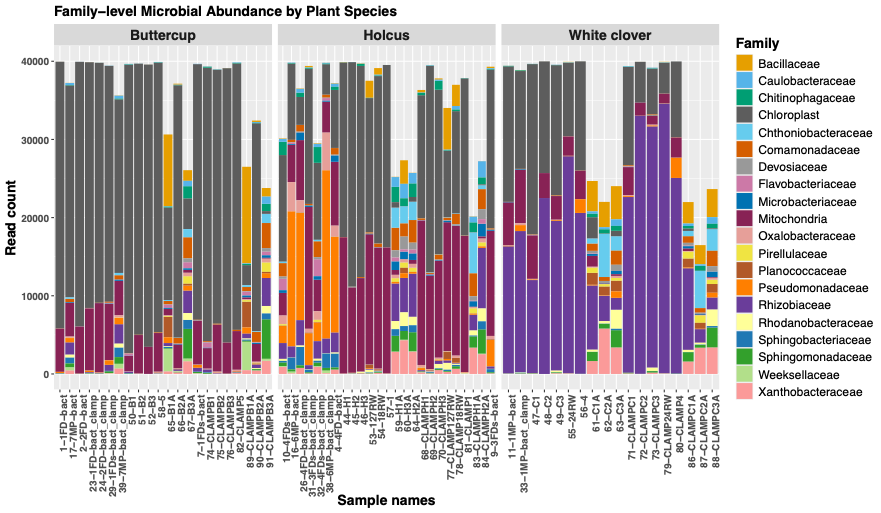

Supplemental Figure 6: Schematic summary of PNA clamp effects on 16S rRNA gene sequence composition across plant species and treatments.

This visual schematic summarises the relative abundance of bacterial, chloroplast, and mitochondrial 16S rRNA gene sequences in three plant species — Buttercup, Holcus, and White clover — under three experimental conditions: No clamp, Standard clamp, and Optimised clamp. Each bar represents the average composition within that treatment group for each plant species.

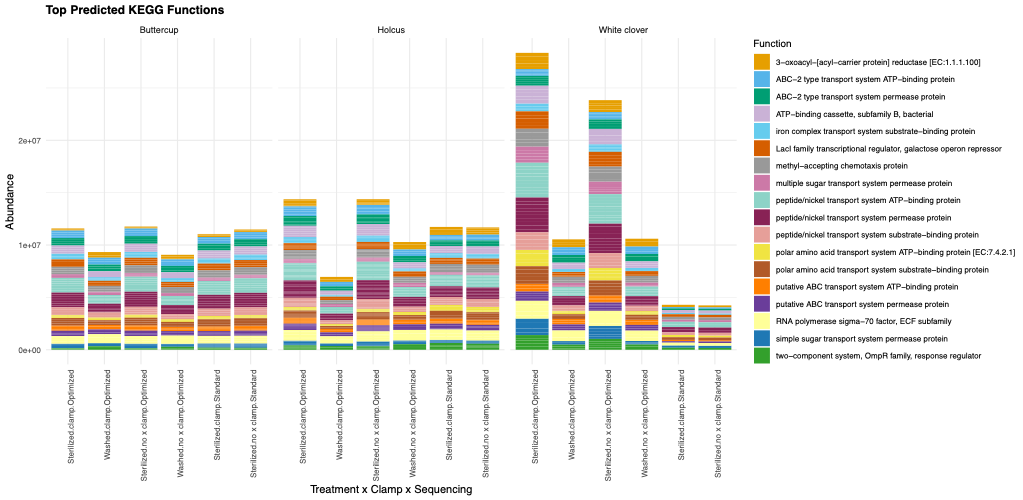

Supplemental Figure 7: Effect of root sterilisation on the discovery of endophytic bacteria and the abundance of predicted functional genes across plant species. Relative abundance of top predicted KEGG functions based on 16S rRNA sequencing from roots of Buttercup, Holcus, and White clover subjected to different treatments (sterilised with clamp, sterilised without clamp, and washed control

**Table 1. PERMANOVA for endophytic bacterial community composition**

| **Factor** | **Df** | **Sum of squares** | **R²** | **F** | **p-value** |
| --- | --- | --- | --- | --- | --- |
| **Treatment** | 2 | 54,649 | 0.216 | 8.564 | 0.001 |
| Residual | 62 | 197,827 | 0.784 | — | — |
| Total | 64 | 252,475 | 1.000 | — | — |
| **Plant species** | 2 | 16,936 | 0.067 | 2.229 | 0.004 |
| Residual | 62 | 235,539 | 0.933 | — | — |
| Total | 64 | 252,475 | 1.000 | — | — |
| **Clamp status** | 1 | 1,406 | 0.006 | 0.353 | 1.000 |
| Residual | 63 | 251,069 | 0.994 | — | — |
| Total | 64 | 252,475 | 1.000 | — | — |

PERMANOVA was performed on centred log-ratio (CLR)–transformed ASV tables using Euclidean distances with 999 permutations.
